# Genetic Modifiers Influencing the Acute and Long-Term Responses to Traumatic Brain injury in *Drosophila*

**DOI:** 10.1101/2025.02.10.637558

**Authors:** Brandon Molina, Jesse J. Rojas, Nadja El-Mecharrafie, Eric P. Ratliff, Marta M. Lipinski, Kim D. Finley

**Affiliations:** Donald P. Shiley BioScience Center, San Diego State University, San Diego, CA, USA; Department of Biology, San Diego State University, San Diego, CA, USA; Shock, Trauma, and Anesthesiology Research (STAR) Center; Department of Anesthesiology, University of Maryland School of Medicine, Baltimore, MD, USA

## Abstract

Worldwide, traumatic brain injury (**TBI**) represents a major cause of mortality and long-term disability, with even mild repetitive forms of TBI (**mTBI**) potentially having deleterious consequences for both the acute and long-term function of the nervous system. Understanding the key cellular and molecular processes that occur following TBI exposure and potential factors that influence individual injury responses has been limited in part by the lack of comprehensive *in vivo* screening technique that directly compares stress sensitive and resistant genotypes involving conserved genes and functional pathways. In this report, we use a high throughput adult *Drosophila* trauma model system to compare the impact modest insulin signaling (IRS/c*hico*) and autophagic defects (*Atg8a/MAPLC3, Ref(2)P/SQSM1)* have on traumatic injury outcomes. Using both severe and mild repetitive injury conditions the acute mortality indexes, longevity profiles, molecular and behavioral changes (locomotor, sleep) for individual fly genotypes were assessed. Compared to control cohorts (*w^1118^*/+), heterozygous *chico* mutants (*chico^1^/+*) demonstrated resistance, while aged flies (+2-weeks) or autophagy mutants (*Atg8a^1^*, *Ref(2)P^c/e^*, *Ref(2)P^e/+^*) showed heightened trauma sensitivity. Alterations that promote or impair autophagic function, longevity and stress responses were examined. Overall, this injury paradigm illustrates the effectiveness of using model systems to characterize conserved genetic factors that influence neuronal autophagy during complex trauma response. It also raises the potential for developing unique screens and therapies for patients that have experienced TBI.

## INTRODUCTION

Traumatic brain injury (**TBI**) remains a major cause of mortality and long-term neurological disability throughout the world. In addition to severe injuries there has been a growing interest in the long-term health problems associated with low-impact, multi-bout types of head injuries, which are often associated with military service, contact sports or domestic violence. While significant strides have been made on the assessment and evaluation of TBI and concussion severity, at this time treatment options remain somewhat limited and largely involve monitoring and supportive care. Therefore, a better understanding of potential risk factors may be helpful in gaging potential outcomes and assist with the design of customized therapies. Broadly speaking, components that influence TBI responses can be broken down into external (medical care access and pre-existing health conditions) and intrinsic (age, gender, genetics) risk factors, which together with the type and extent of trauma could have a major impact on an individual’s outcome [1-4]. Additional factors also influencing outcomes are highlighted by the complex interplay that occurs between multiple cell types (neurons, glia) and signaling cascades that regulate appropriate activation of inflammation, clearance, and repair pathways [5-8]. Delays in healing or secondary rounds of damage linked to protracted inflammatory responses, may tip the balance toward poor outcomes [9-12].

Detailed genetic studies into trauma-related risk factors have partly been hampered by the limited high throughput *in vivo* models, further hindered the development of effective diagnoses and treatments [13,14]. One of the most striking intrinsic factors influencing vastly different outcomes involves an individual’s age at the time of trauma. While children and young adults are disproportionately exposed to TBI, people in older demographics often show a heightened sensitivity to head trauma and relatively poor long-term outcomes [2,7,12,15,16]. Age-related differences likely involve individual differences to pre-injury cognitive reserve and extent of pre-symptomatic neural degenerative disease as well as baseline changes or progressive defects in regulating neural inflammation, signaling or repair mechanisms. Targeted human genome-wide association (**GWAS**) based studies have provided useful insight, despite the difficulties working with the complexity of head trauma and the relatively limited number of individuals for a given study [17-19]. Research has initially focused on known neurological factors including allelic variants of the Apolipoprotien-E (APOE) gene, which has also been associated as a risk factor for the development of Alzheimer’s disease (**AD**) [20,21]. GWAS based studies have consistently indicated that carriers with the APOEe4 allele have relatively poor outcomes following different types of TBI exposures [4,22,23]. Other human SNP-based studies have examined links between TBI outcomes and variants of mitochondrial, BDNF and neurotransmitters genes. While intriguing, definitive assessments were not possible due to small sample sizes in the complexity of the analyses [4,23]. These findings indicate that using a high throughput Drosophila model system to identifying conserved genes or allelic variants impacting acute and long-term TBI outcomes, could be used as a powerful tool to identify key components influencing the recovery patterns following head trauma (genetic prescreening) as well as the development of tailored therapeutic treatments.

Previously, we have described a high throughput model of traumatic injury, which inflicts controlled cortical impact-like (**CCI**) conditions on adult *Drosophila* [13,24]. Consistent with human patients and TBI models using rodents, adult flies demonstrate dose dependent phenotypes following the exposure to both severe (sTBI) and mild repetitive (mTBI) conditions. This included the dose-dependent shortening of adult lifespans, subtle behavioral and sleep changes, and the profound activation of inflammation and autophagic pathways [13,24]. Based on this and other studies on metabolic factors that modulate neural degeneration, these studies examine the impact that aging and select defects in autophagic and insulin-signaling components have on *in vivo* trauma responses. Customization of our original *Drosophila* TBI techniques to permit detailed genetic analysis of trauma responses. As an initial proof of principle evaluation, mutant genotypes previously classified as being long-lived and stress resistance (IRS, *chico*) or short-lived and stress sensitive (MAP-LC3, *Atg8a*; SQSTM1, *Ref(2)P*) were assessed [25-28]. Global changes to lifespan profiles, autophagic and inflammation responses as well as differences to locomotor and sleep behaviors were examined as part of the TBI testing platform [13,29,30]. Taken together these studies show that older adult *Drosophila* and flies with autophagic defects (*Atg8a, Ref(2)P*) are trauma sensitive, while fly cohorts with modest insulin signaling defects (*chico*) demonstrate TBI resistance. Using transgenic reagents to target tissue specific gene suppression or enhancement the impact of *Ref(2)P* following TBI was further examined [27,28]. This integrated TBI assessment platform can now be used to quickly identify and assess potential genetic factors or novel therapeutic interventions that promote or inhibit the appropriate neuronal healing that needs to occur to promote not only patient survival rates but optimal recovery following exposure traumatic brain injuries.

## Methods and Materials

### Drosophila Stocks and Culturing Conditions

The Canton-S, *w^1118^*, *Atg8a^1^*, *chico^1^*, *Actin-Gal4, Elav-Gal4*, *Repo-Gal4*, *Elav-Gal4;UAS-Dcr2* stock lines have been described previously and obtained from the Bloomington Stock center (https://bdsc.indiana.edu/) [27]. The *ref(2)P^c03993^* and *ref(2)P^e00482^*stock lines were obtained from the Harvard Medical School Exelixis Stock Collection and are described (http://flybase.org) [22]. The *UAS-ref(2)Pi* (v108193, P[KK105338]) and *UAS-Atg8ai* (v109654, PKK102155]) fly stocks were obtained from the VDRC stock center (https://stockcenter.vdrc.at) and the *UAS-ref(2)P* line was a gift from Tom Neufeld [13,29,30]. For most studies F1 offspring were generated by crossing Canton-S (CS) females with *w^1118^* (WT, *w^1118^*/+), *chico^1^*, *Ref(2)P^c03993^* and Ref(2)P^e00482^ male flies [25,26,31]. Male and female flies were collected and aged in same-sex cohorts (25 flies per vial) and maintained on standard lab media and conditions at 25°C in 55-65% humidity and using a standard 12 h:12 hour light:dark cycle (**LD**) [13,25,29]. For fasting studies, flies were placed in vials containing 1% agarose for 8-hrs (9:00am to 5:00pm) before being collected [31,32].

### Traumatic Brain Injury

Flies were incapacitated, placed in a clean 2-ml screw cap tube (10 flies/tube) and allowed to recover before being traumatized [13]. For injury, tubes containing flies were placed into the Omni Bead Ruptor-24 homogenizer (Omni International, Kennesaw, GA, USA) and subjected to specific pre-programmed shaking conditions that have been previously described [13]. A severe trauma bout (sTBI) was set at 4.3 m/s and a mild bout (mTBI) at 2.1 m/s. For multi-bout conditions, fly cohorts were injured (5 seconds) and allowed to recover for 30 seconds before the start of subsequent injury bouts. Following the completion of the injury, all fly cohorts were placed into vials, which were placed on their sides to allow for recovery [13].

### Mortality and Longevity Profiles

The 24-hr mortality indexes (24MI) were determined by subjecting for different fly cohorts to a single 5-second injury sTBI bout and determining the percent of dead flies after 24 hours [13,29,30]. The longevity profiles of flies were determined after subjecting fly cohorts (n = +100) to a single sTBI or mTBI (10x) conditions at 1 or 3-weeks of age and the number of dead flies were recorded 3 times each week [13,29].

### *Imaging* Studies

For immunofluorescence studies an average of 12 to 14 heads from 1-week old WT (*w^1118^*/+) or *Ref(2)Pc/e* null female flies were collected from untreated (controls) or groups that experienced fasting (8-hr) or mTBI 10x (24-hr) conditions. Brains were immediately dissected, fixed in 4% paraformaldehyde, 1xPBS (4% PFA-PBS) and stained overnight (4C) with anti-Elav (1:500 dilution, Cat. No. 9F8A9, Developmental Studies Hybridoma Bank [DSHB], Iowa City, Iowa, USA), and anti-Atg8a antibodies (1:250 dilution, anti-GABARAP, E1J4E, Cell Signaling Technology [CST], Danvers, MA, USA) [13]. Washed tissues were blocked (5% NGS), incubated with Alexa Fluor-488 (1:250 anti-mouse) or Cy3 (1:250 anti-rabbit) secondary antibodies (Jackson ImmunoReseach Labs, Inc.) and mounted using Vectashield (Vector Labs, Burlingame, CA, USA) as previously described [13,33,34]. One-micron images taken at the same magnification, excitation and collection settings were collected from adult fly brains using a Zeiss 720 confocal microscope (SDSU Biology Department Imaging Core). The average number of Atg8a positive puncta from ELAV-positive cortical neuronal cell bodies (n = 30) were determined for each fly group from a series of 10 micron^2^ fields as previously reported [13].

### Western Analysis

Male flies (1-week) from WT (*w^1118^*/+), *Ref^c/e^*, *Ref^e^/+*, *Elav-Gal4*/+ and *Elav-Gal4/UAS-dsRef(2)Pi*/*UAS-dicer2* (iRef(2)P) genotypes were exposed to fed or fasted (8-hrs) conditions.[22,24,25] A second cohort of male flies from WT (*w^1118^*/+), *chico^1^/+*, *Ref^e^/+*, *Ref^c/e^* and *Elav-Gal4/UAS-dsRef(2)Pi*/*UAS-dicer2* (Neural-iAtg8a) genetic backgrounds were exposed to control or 10x mTBI (2.1 m/s) conditions [13]. At defined timepoints flies were collected, flash frozen and stored (-80°C) [33]. Separated heads were extracted with lysis buffer (2% SDS, 150mM NaCl, 50mM Tris, pH 7.5) and protease inhibitors (Thermo Scientific/Pierce, Rockford, IL, USA) using the Bead Ruptor-24 System (Omni International, USA). Cellular debris was removed (10,000 x *g*, 10 min) and the protein concentrations of supernatants were determined using the DC Protein assay (Bio-Rad) using established protocols [33]. Protein samples (20 μg) were resolved on a 12% Bis-Tris gel (Bio-Rad) and transferred onto Immobilon-P PVDF membranes (Millipore Corp., Billerica, MA, USA) using the Trans-Blot Turbo system (Bio-Rad). Each blot was sequentially probed for Atg8a-I/II (1:500 dilution, E1J4E, CST), Ref(2)P (1:1,000 dilution), Ubiquitin (1:1,000 dilution, P4D1, CST) and Actin (1:2,000 dilution, JLA20, DSHB) proteins and digital Imaging System and Quantity One software (Bio-Rad) and established reagents and protocols [13,31].

### Quantitative PCR

Control and mTBI (10x) treated male flies (1-week) from WT (*w^1118^*/+), *chico^1^/+*, *Ref^e^/+* genotypes were collected at 0-hr, 4-hr and 24-hr time-points, flash frozen and stored (-80°C). RNA was isolated from 25 heads using Trizol (ThermoFisher Scientific, Grand Island, NY, USA) and cDNA libraries generated using the RevertAid First Strand cDNA Synthesis kit, with a combination of random hexamer and oligo-dT primers (Thermo Scientific, Pittsburg, PA, USA) [29,30,32]. Quantitative PCR was performed on a CFX Connect Real-Time PCR Detection System (Bio-Rad) and Universal PCR SYBR Mix reagents (Bio-Rad). Primer sequences for the *AttC*, *DptB*, and *Mtk Drosophila* genes are available upon request. Melt curve analyses of all qPCR products confirmed the production of a single DNA duplex. The Pfaffl method was used to quantitate expression profiles and *Exba* (*Xba*) was used as the reference gene. Relative mRNA levels of non-injured flies were set at 1.0 and subsequent expression levels from different time points expressed as normalized values [29,30,32].

### Negative Geotaxis response

The rapid iterative negative geotaxis (**RING**) protocol and apparatus has been described previously [25,29,30]. Digital images of climbing behaviors were taken for fly cohort (n ≥ 100) before and 5-days following mTBI 10x exposure. Replicate bumps (n = 3), with a 1-min rest between tests, were conducted to assess average distance traveled (cm) after 5 seconds. Digital images were used to score the distance traveled from 0 (bottom) to 6 cm (top) of individual flies [25,29,30]. Triplicate runs were used to establish average climbing indexes, SEMs and 25% or greater decline in climbing behaviors between young (1-week) and aged (3-weeks) control and mTBI fly cohorts [25,29,30].

### Sleep-Circadian Analysis

Male flies (1-week) for each genotype were subjected to 1x, 5x or 10x mTBI bouts (2.1 m/s) and allowed to recover under standard 12hr:12hr LD cycle conditions for 5 days. Individual flies were transferred to tubes (5 mm x 65 mm) containing fly media, placed into a DAM5 System (Trikinetics Inc., Waltham, MA, USA) and allowed to acclimate for 2-days under 12hr:12hr LD cycle conditions [25,29]. Fly cohorts were placed in constant dark conditions (**DD**) and movement (beam breaks) collected in1-minute bins for 6-days using DAM system v3.8 software and used to generate individual actograms [25,29]. Data sets were further analyzed using the Sleep-Lab software (MatLab) developed by Dr. William Joiner, as previously described [25,29,35]. Any flies that died during the sleep study periods were removed from analysis. The waking activity, brief awakenings, daily sleep bouts, and sleep bout duration during CT0-12 and CT12-24 time periods were averaged over a 5-day period and calculated using the Sleep-Lab software [25,26,31]. Brief awakenings refer to 5-min periods with four or fewer activity counts that were preceded and succeeded by a minimum of 5 minutes of inactivity [36].

### Statistical Analysis

Data are presented as group averages and ± standard error of the mean (SEM) were calculated using Excel. Statistical analysis between control and individual experimental groups was performed using Student’s T-test (two-tailed, unpaired) to establish significance is indicated as \**p* < 0.05, \*\**p* < 0.01, and \*\*\**p* < 0.001 (GraphPad Prism 6, https://www.graphpad.com) [29,35].

## RESULTS

### sTBI mortality indexes (MI^24hr^) and Lifespans of traumatized Drosophila

Previously, we established an adult *Drosophila* model that delivers highly reproducible levels of severe (sTBI) and mild repetitive bouts of traumatic injury (mTBI) to large numbers of flies. This platform involved using the Omni Bead Ruptor-24 Tissue Homogenizer, which can be programed to deliver different intensity (meters/second, [m/s]), duration or number of injury bouts to adult flies (10 per vial) [13]. For initial sTBI studies, wildtype (WT, *w^1118^/+*) male and female flies at 1, 2, 3 and 5-weeks of age were exposed to a single sTBI bout (1x, 4.35 m/s) and the number of dead flies counted 24-hrs following trauma. At 1-week, young adult flies showed the lowest mortality level (∼12%), while 5-week-old fly cohorts demonstrated the highest MI^24hr^ profiles (93% and 74.5%, see **Table 1**) [13]. The responses of young flies from different genetic backgrounds to sTBI were also examined. Previously, we had shown that modest insulin signaling defects promoted longevity and the stress responses of adult Drosophila. For this study, the MI^24hr^ profiles of 2-week-old *chico^1^/+* heterozygous male and female flies (insulin receptor substrate, IRS) were significantly lower than age-matched controls (2.3%, 1.4%, **Table 1**) [31]. Short-lived mutants that corresponded to the human *MAP-LC3* (*Atg8a*) and *p62*/*SQSTM1* (*Ref(2)P*) genes were also examined. At 2-weeks of age, *Atg8a^1^* male and female flies showed heightened sensitivity to sTBI conditions (82%, 44%), as did 1-week old *Ref(2)P^c/e^* null (73%, 53%) and heterozygous fly cohort controls (45% and 53%, see **Table 1**) [31]. Also examined were the in MI^24hr^ responses of flies with mutation in the *Drosophila Ref(2)P* gene, which is the mammalian *p62* (*SQSTM1*) homologue [26]. Overall, male flies from any age or genetic background had higher MI^24hr^ profiles than matching female cohorts. Individual fly cohorts were exposed to mTBI (10X) conditions at 1-week of age and used to establish trauma-induced changes to longevity [13].

**Table 1.**
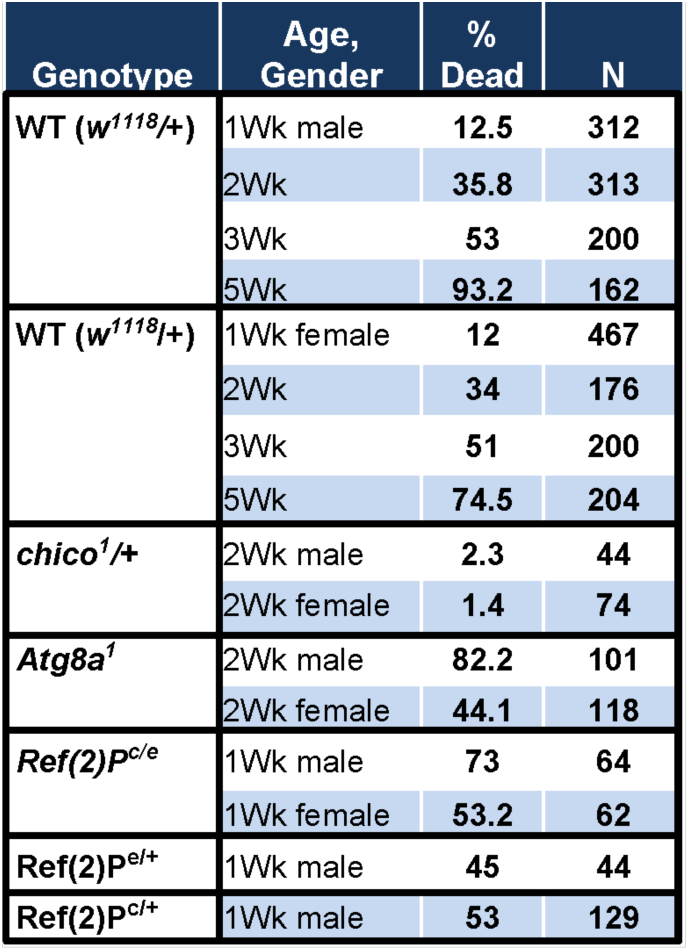
24-hr sTBI Mortality Indexes (4.35 m/s)

Previously, we have established genotype specific lifespan profiles, which are illustrated in **Table 2**. Exposed to mild repetitive TBI conditions (mTBI, 10x bouts) each cohort of flies showed a significant reduction to average (days) or MI-50% (days) lifespan profiles [13]. Comparing the percent change between control and mTBI exposed cohorts for each genotype determined that homozygous *Atg8a^1^* and *Ref(2)P^e/c^*mutants showed an accelerated MI-50% response (days) (**Table 2**). Consistent with the global age-dependent increase in trauma sensitivity, flies from short-lived genotypes showed elevated sensitivity, while long-lived strains were partly resistant following TBI exposure

**Table 2.** Average Lifespans with mTBI (10x)

| Genotype | mTBI | Ave (days) | % Change | N |
| --- | --- | --- | --- | --- |
| WT | 0 | 46.9 |  | 382 |
|  | 10x | 26.3 | -43.9 | 285 |
| *chico<sup>1</sup>/+* | 0 | 61.4 |  | 186 |
|  | 10x | 43.1 | -29.8 | 170 |
| *Ref(2)<sup>P<sup>c/e</sup></sup>* | 0 | 37.4 |  | 77 |
|  | 10x | 19.3 | -48.4 | 104 |

### Autophagic Responses of Adult Ref(2)P Mutants, Confocal Analysis

To examine the sensitivity that Ref(2)P homozygous and heterozygous mutants to trauma required clarification of autophagic responses in the adult nervous system. Previously, we had examined the impact that fasting had on autophagy in the adult nervous system of WT, *Atg8a* and *chico* mutant flies [24]. In addition, we demonstrated that exposure to mTBI (10X) leads to dynamic acute and long-term changes to the pathway in WT nervous system [13]. Therefore, we established the autophagic profiles that occurs within the adult Ref(2)P mutant nervous system. Initial assessments of autophagic responses involved imaging the distribution of Atg8a-positive puncta or autophagosomes throughout the CNS tissues from adult WT or *Ref(2)P^e/c^* Drosophila [13]. Flies were left untreated or exposed to fasting (8-hr) or mTBI (10X, 24-hr post) conditions before the brains (∼12 per condition) were collected and immuno-stained using anti-Atg8a protein (red) and anti-Elav antibodies (green, **Fig. 1A**) [13]. Serial confocal images were obtained for each brain and the number of Atg8a positive puncta (red) counted and averaged for multiple fields (10 um^2^) within cortical regions that contained neural soma (Elav+, green, **Figure 1A**) [13,37]. Consistent with previous findings, control flies showed an increase in the average number of Atg8a+ puncta (10 μm^2^) in neurons following a fast or mTBI exposure (**Figure 1A-C**). The average number of Atg8a-positive puncta was consistently lower in the neural tissues from *Ref(2)P^e/c^* under basal conditions (**Figure 1B, 1D**). Fasting did not alter the average number of Atg8a+ puncta and TBI exposure produced a modest increase in the number of autophagosomes in *Ref(2)P^e/c^* mutant neurons (**Figure 1B**) [26].

**Figure 1.**
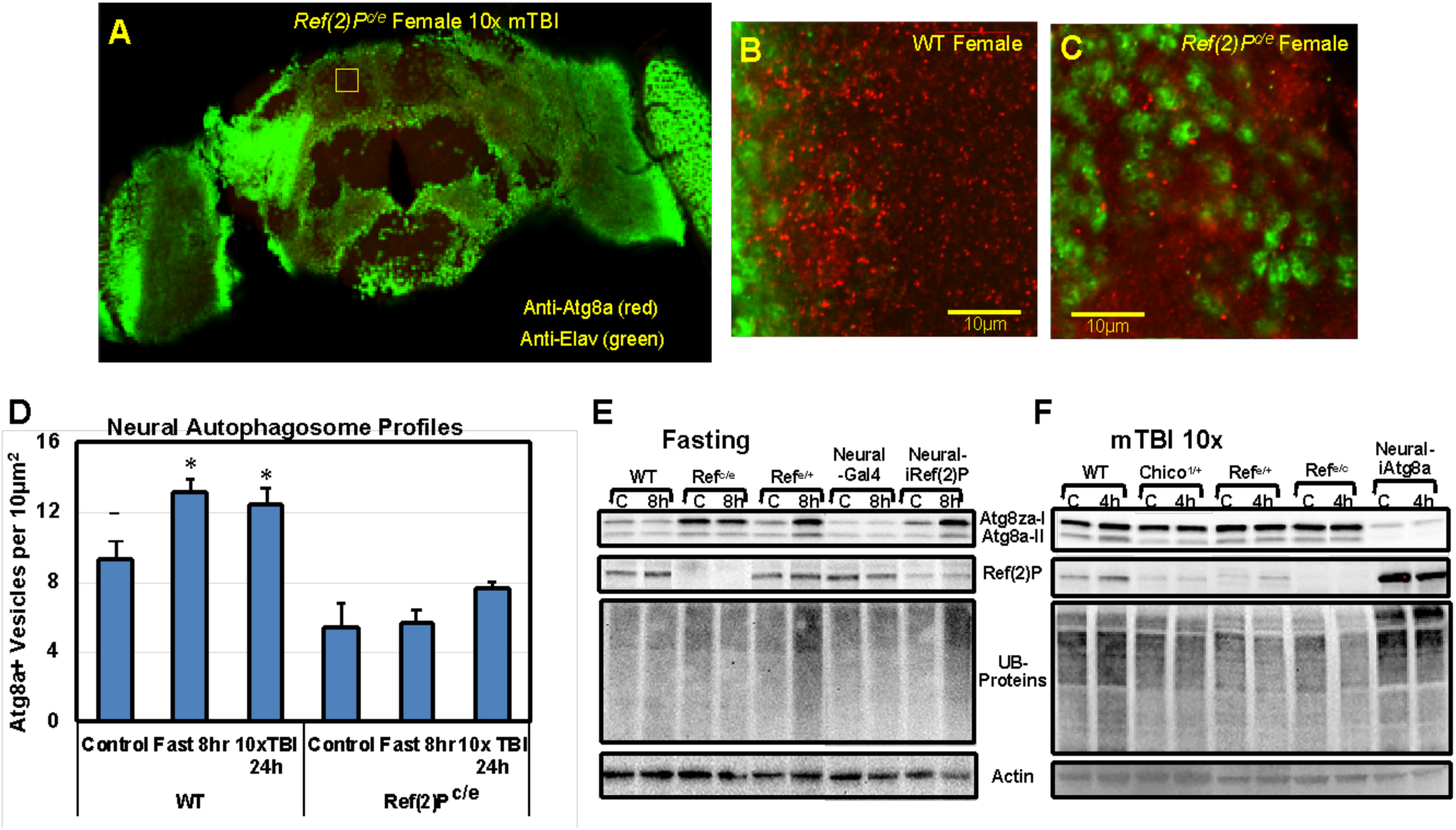
Genetics and Adult Drosophila Neuronal Autophagic Responses to Fasting and mTBI. **A-C**) Representative and enlarged neural images (highlighted inset) of Atg8a staining patterns. Brains were co-stained for neuronal nuclei (anti-Elav; green) and autophagosomes (anti-Atg8a; red). **B-C**) Neural soma from WT (n=12) and *Ref(2)P* (n=14) nulls 24-hours following mTBI exposure(10x, 2.1 m/s). **D**) Autophagsomes in control, fasted (8-hours) and traumatized (24-hours) WT and *Ref(2)P* neural tissues (24 fields each, 10 um^2^ per condition, P value *<0.05). **E-F**) Western blot analysis of adult *Drosophila* neural tissues (heads) sequentially probed for Atg8a-I/II, Ref(2)P, Ubiquitin and Actin proteins. **E**) Neural profiles from fed and fasted (8-hrs) WT, *Ref^c/e^*, *Ref^e^/+*, *Elav-Gal4*/+ and *Elav-Gal4/UAS-dsRef(2)Pi*/*UAS-dicer2* (iRef(2)P) fly cohorts and blots **F**) Westerns were prepared from control and traumatized (4-hour post mTBI) fly cohorts from WT, *chico^1^/+*, *Ref^e^/+*, *Ref^c/e^*and *Elav-Gal4/UAS-dsRef(2)Pi*/*UAS-dicer2* (Neural-iAtg8a) genetic backgrounds. P value *≤ 0.05.

### Western analysis

Western blot analyses has been used to examine the autophagy responses of cells and tissue by examining changes to Atg8a-I and Atg8a-II (LC3-I/LC3-II) ratios and Ref(2)P (p62) and ubiquitinated protein levels [25,26,31]. Our work and that of others have shown the pathway is transiently activated following exposure to multiple stressors. For this study young (1-week) male WT (*w^1118^/+*), *Ref(2)P^e/c^*, *Ref(2)P^e/+^*, *Elav-Gal4*/+ (neural) and *Elav-Gal4/UAS-dsRef(2)Pi* (neural KD) cohorts were exposed to standard/control (**C**) or fasting conditions (8-hr), collected, flash frozen and total protein extracts prepared from head tissues. Western blots were prepared from head extracts and sequentially probed for Atg8a, Ref(2)P and ubiquitinated proteins (**UB-proteins**) and actin (**Figure 1E**). As seen previously, there are distinct genotype-specific differences in basal autophagic profiles (**Figure 1E)** [24]. This included the elevated Atg8a-I and Atg8a-II levels in control *Ref(2)P^e/c^* null flies, which was consistent with imaging studies and further indicates impaired autophagosome formation (**Figure 1E**) [9,24]. Following an 8-hour fast *Ref(2)P^e/c^* nulls showed minimal changes to Atg8a-I and Atg8a-II profiles in Drosophila neural tissues, while fasting revealed significant autophagic defects in LOF Ref(2)P genetic backgrounds (*Ref(2)P^e/+^* and neural-*dsRef(2)Pi*, **Figure 1E**) [13,25,31]. Directly examining Ref(2)P levels underscored genotype specific protein levels and the buildup of UB-proteins highlighted inability of autophagy to clear targeted substrates in *Ref(2)P^e/+^* and *iRef(2)P-KD* flies (**Figure 1E**) [13,25,31].

Western analysis is also used to examine acute neural autophagic responses following mTBI (10x) (**Figure 1E-F).** Young (1-week) control WT (*w^1118^/+*), *chico^1^/+*, *Ref(2)P^e/+^*, *Ref(2)P^e/c^*, and *Elav-Gal4/UAS-dsAtg8ai* (neural KD) and mTBI (10x) treated male fly cohorts (4-hrs post) were collected, flash frozen and total protein extracts from heads used for Western blot analysis of Atg8a-I, Atg8a-II, Ref(2)P, UB-protein and actin profiles (**Figure 1F**) [13,25,31]. The acute buildup of Atg8a-I, Atg8a-II, Ref(2)P, UB-protein levels for WT neural tissues 4-hrs after mTBI (10x) exposure, flies demonstrated acute pathway activation that precedes the dynamic trafficking, fusion and elimination of targeted substrates by lysosomes at later time points (**Figure 1F**) [33]. Trauma resistant *chico^1^/+* mutants, which start with elevated autophagy rates in the CNS (low Atg8a-II levels) also showed an increase in Atg8a-I/II the amounts, indicating the pathway was further activated. In contrast, flies with impaired autophagy (*Ref(2)P^e/+^*, *Ref(2)P^e/c^*, neural*-dsAtg8ai*) demonstrated minimal changes to targeted substrate levels following mTBI exposure indicating a significant delay (**Figure 1F**) [13]. Together, the rapid activation of autophagy in WT and *chico^1^/+* flies and impaired responses in pathway mutants are consistent with global resistance and sensitivity profiles for individual genotypes.

### Inflammatory Responses

An additional molecular system that demonstrates dynamic activation following TBI exposure in both mammalian and *Drosophila* in damaged tissue models, involves the innate immune and inflammatory pathways [13,29,38,39] Under a wide range of conditions inflammation signaling plays a critical role in communicating damage, which in turn leads to the further recruitment of immune components and activation of downstream cellular repair processes within effected tissues [6,40,41]. Additional studies indicate the insulin and autophagy pathway activity can influence both the basal and acute regulation of inflammatory components [25,31]. In flies, the innate immune systems are largely analogous the mammalian Toll (Toll-like receptor) and the IMD pathways (Tumor necrosis factor receptor) [42-44]. Activation of these pathways directly impacts the regulation of *Drosophila* anti-microbial peptides (**AMP**), including *AttC*, *DptB,* and *Mtk* that also have cytokine/chemokine-like functions in flies. [45-47]. Previously, TBI exposure has been shown to result in a robust increase in *AttC*, *DptB,* and *Mtk* mRNA levels in the adult fly CNS. Typically, we find that following exposure to mTBI (10x) conditions there is a rapid peak of expression at 4-hrs, which quickly returns to near baseline levels after 24-hrs [13,46]. To determine whether different genetic backgrounds would also influence inflammation, WT (*w^1118^/+*), *chico^1^/+* and *Ref(2)P^e^/+* male flies (1-week) were exposed to standardized mTBI (10x) conditions [22,24]. Triplicate cohorts were collected at 0 (control), 4-hr and 24-hr time points, RNA isolated from heads and replicate cDNA libraries generated for each condition [26,28]. The expression levels of *AttC*, *DptB*, and *Mtk* transcripts were determined using gene specific primers and qRT-PCR analyses (normalized to *Cyp1*). Basal levels of each AMP in neural tissues from each genotype are illustrated in **Figure 2A**.

**Figure 2.**
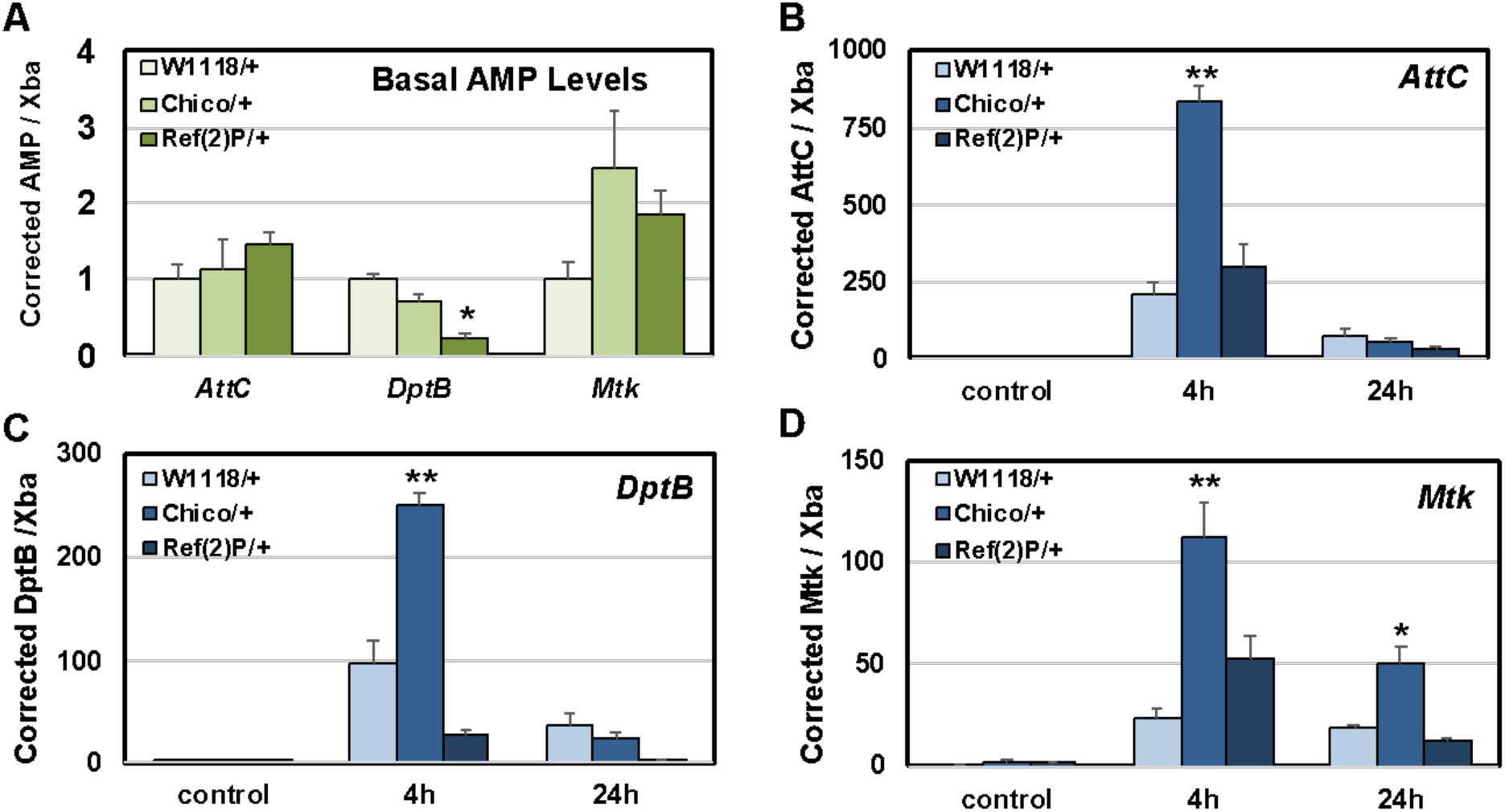
Genotype Specific Inflammatory Responses in Neural Tissues Following mTBI (10x). Young male WT (*w^1118^*/+), *chico^1^*/+, and *Ref(2)P^e^/+* heterozygous flies (1-week) were exposed to mTBI. Neural cDNA libraries (3x) were generated for qRT-PCR analyses from each genotype and 0-hr, 4-hr and 24-hr time points and the expression levels of antimicrobial peptides (AMPs) determined. **A**) Basal *AttC*, *DptB*, and *Mtk* expression levels in **B**) *AttC*, **C**) *DptB* and **D**) *Mtk* trauma (mTBI, 10x) related expression changes in WT, *chico^1^*/+, and *Ref(2)P^e^/+*. All values were corrected using *Exba* gene expression as the loading control. P values *≤ 0.05; **≤ 0.01.

When compared to WT flies both the *chico^1^/+* and *Ref(2)P^e^/+* genotypes showed unique basal AMP profiles, with DptB and Mtk expression showing substantial differences (**Figure 2A**). Following TBI exposure all three fly genotypes showed the normal dramatic increase in AMP expression levels (4-hr), followed by a rapid recovery (24-hr, **Figure 2B-D**). Overall, *chico1/+* mutants showed a highly significant increase in expression profiles for all three AMPs, with a rapid return to near baseline levels for AttC and DptB transcripts (**Figure 2B-D**) [31,48]. The exception was Mtk, which was significantly higher at 24-hrs post-TBI than that of matching WT controls (**Figure 2D**). The *Ref(2)P^e^/+* flies also showed unique expression profiles following trauma, with *AttC* demonstrating nearly WT patterns, *DptB* having dampened profiles, and *Mtk* showing an accentuated response 4-hrs post-TBI (**Figure 2B-C**) [26]. Together with altered mortality profiles and autophagic responses, this study indicates there are genotype specific inflammatory responses that reflect resistance or sensitivity to trauma. In addition, the unique basal and TBI-dependent expression differences detected for *Mtk* could in part reflect its regulation by both the Toll and IMD pathways [38].

### Behavioral Changes as Markers of Trauma Resistance or Sensitivity

To clarify the impact that mild repetitive trauma exposure has on adult Drosophila nervous system, we had examined two robust adult Drosophila behaviors that include the negative geotaxis response (**NGR**) and circadian sleep-related profiles [13,25,29]. Both the NGR and sleep-based assays are robust techniques that have been used to detect even minor functional changes to the CNS, which we had demonstrated was altered by TBI exposure. This suggested that both behaviors could be used as part of a larger testing platform to evaluate the influence that genetic factors have on trauma responses. Therefore, following minor modifications, flies from different ages or genetic combinations were assessed for TBI-dependent changes to behaviors that in turn reflect neurological defects.

### Negative Geotaxis Responses

We have previously detailed the NGR protocol and shown that as flies age, they show a highly reproducible decline in climbing abilities that can be influenced by therapeutic, genetic and dietary factors [25,30,31,49]. Other mTBI-based studies showed only a modest impact to climbing behaviors shortly after trauma. The assay was modified for this study, where baseline NGR levels were obtained for young (1-week) or middle-aged (3-weeks) female fly cohorts (∼50 each), before being exposed to mTBI (10x) conditions and allowed to recover [13]. The climbing behaviors were then assayed 5-days post trauma, thereby each cohort of flies served as its own control. As seen previously, young WT flies (1-week) showed a modest reduction in NGR profiles 2-days following trauma, while WT middle-aged (3-week) cohorts showed a pronounced decline, indicating a heightened sensitivity to trauma (**Figure 3A**). The geotaxis profiles of matched *chico^1^/+* fly cohorts were also examined and showed similar age and trauma-related changes, with the exception that aging has less of an impact on geotaxis profiles following mTBI exposure (**Figure 3A**). Therefore, a decline in geotaxis responses that was 25% or greater, 2-days following mTBI exposure was set as the trauma sensitivity cutoff. The NGRs of *Ref(2)P^e/c^* and *Atg8a^1^* mutant flies were also examined. Before trauma, young flies from both genotypes had relatively normal geotaxis profiles as well as the natural age-dependent decline for this behavior. However, using this assay young *Ref(2)P^e/c^* and *Atg8a^1^* flies (1-week) also showed a marked sensitivity to trauma that was further exacerbated by age (**Figure 3A**). Female fly cohorts from the same genotypes and ages (1 or 3-weeks) were also examined and showed nearly patterns of sensitivity or resistance (see **Supplemental Figure S1**).

**Figure 3.**
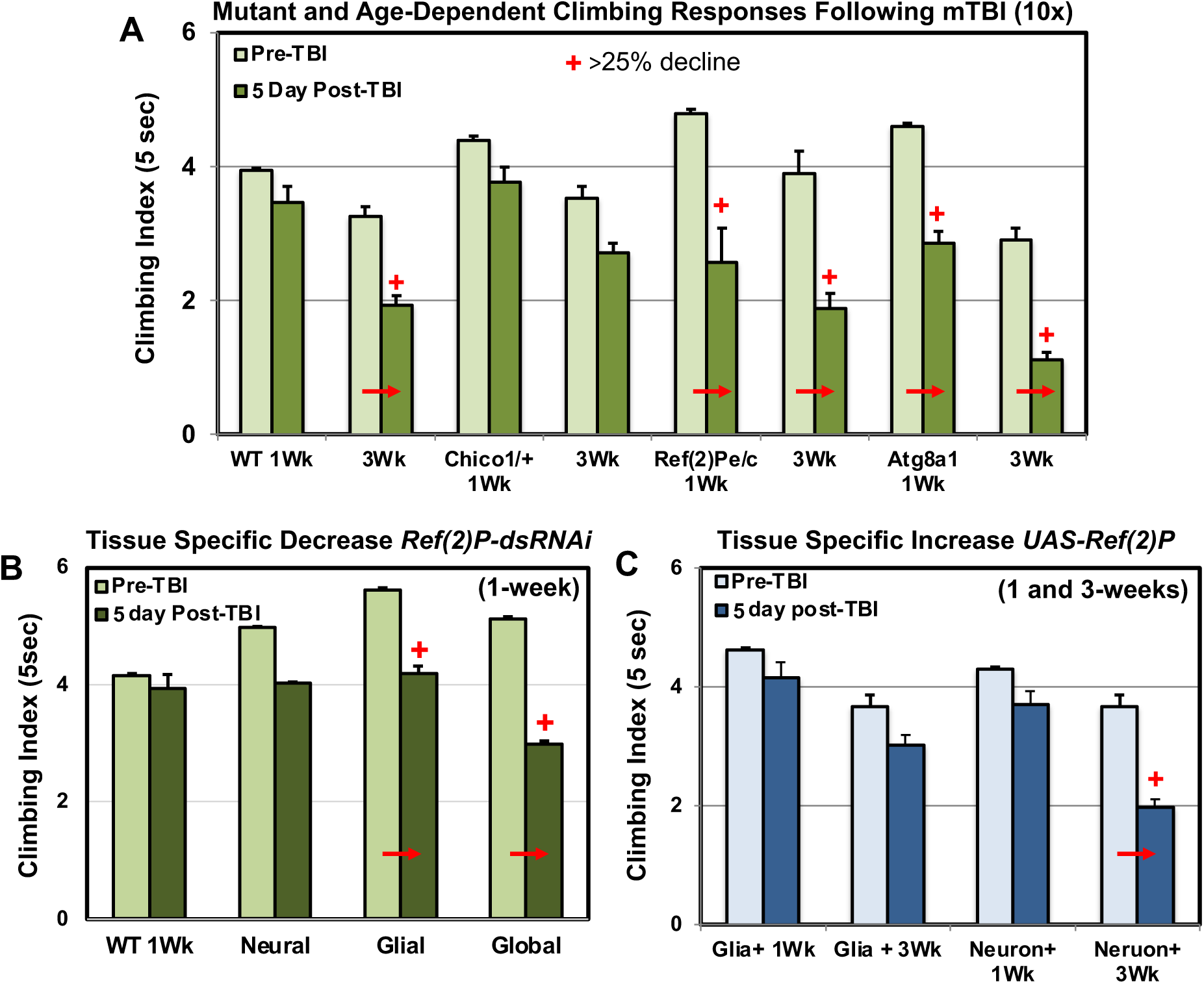
Genotype and Age-Related Changes to Climbing Behaviors (NGR) of Male Flies Following mTBI (10x). Behavioral changes that are greater than 25% before and following mTBI exposure are indicated (**+**). **A**) The NGR profiles of 1 and 3-week old WT (*w^1118^*/+), *chico^1^*/+, *Ref(2)P^e^/^c^* null and *Atg8a^1^* mutant male fly cohorts before and 5-days following mTBI (10x) exposure. See Supplemental data for matching female NGR profiles (Figure S1). **B**) Climbing responses of flies before and after mTBI (10x) exposure that have neural (*Elav-Gal4/UAS-iRef(2)P*), glial (*Repo-Gal4/UAS-iRef(2)P*) or global (*Actin-Gal4/ UAS-dicer2/UAS-iRef(2)P*) knockdown of the *Ref(2)P* message. **C**) Climbing responses of young and aged flies elevated *Ref(2)P* expression in glial (*Repo-Gal4/UAS-Ref(2)P*) or neuronal (*Elav-Gal4/UAS-Ref(2)P*) tissues.

The flexibility of the geotaxis assay permitted more detailed examination of the role that the Ref(2)P protein in neuronal or glial tissues has on whole animal TBI responses. For these assays the Drosophila Gal4/UAS system was used to generate adult flies that had a global (*UAS-dsRef(2)P;Actin-Gal4*), glial (*Repo-Gal4*) or neuronal (*Elav-Gal4;UAS-Dcr2*) specific transgenic knockdown (**KD***, UAS-dsDcr2i*) of the *Ref(2)P* message [25,30]. As with *Ref(2)P* null flies (**Figure 1E-F**) different transgenic KD combinations reducing *Ref(2)P* gene expression produced normal adult flies with robust behavioral (NGR) profiles (**Figure 3B**). Following mTBI (10x) exposure, flies with a global *Ref(2)P* KD showed a sharp behavioral decline in that was nearly identical with that of nulls (**Figure 3A-B**). Following trauma, flies with neuronal or glial *Ref(2)P* KDs also showed a significant decline in climbing abilities, with the *Repo-Gal4;UAS-dsRef(2)Pi* transgenic flies demonstrating a rapid and significant reduction in NGR profiles (<25%, **Figure 3B**).

To determine whether elevated *Ref(2)P* levels could have a positive impact on trauma responses, transgenic flies were also generated Ref(2)P was over-expressed in glial or neuronal cells. Young (1-week) and middle-aged (3-weeks) *Repo-Gal4/UAS-Ref(2)P* and *Elav-Gal4/UAS-Ref(2)P* flies had basal climbing behaviors that were similar to that of age-matched control flies (**Figure 3A-C**). In addition, suppression of the age-dependent trauma sensitivity was also examined. In this case, middle-aged *Elav-Gal4/UAS-Ref(2)P* cohorts showed a sharp decline in climbing abilities, while the NGR of matching 3-week old *Repo-Gal4/UAS-Ref(2)P* flies were in large part preserved (**Figure 3C**). Taken together these sensitive behavioral assays further highlight the potential role the Ref(2)P protein plays in the response of glial cells following traumatic injury. It also suggests that behavioral responses following TBI exposure can be used as part of a screening mechanism to identify genetic and potentially environmental factors that have the potential to alter trauma responses [50,51].

### Genotype Specific Alterations to Sleep-Circadian Profiles Following mTBI

Multiple studies, both in humans and rodent models, have demonstrated an increase in behavioral defects that arise due TBI exposure, which include alterations to circadian locomotor activity as well as problems establishing (insomnia) or maintaining sleep (fragmentation) [52-54]. In our original characterization of trauma, we demonstrated that adult female flies exposed to trauma had significant disturbances to their sleep/wake cycle [13]. Here male fly cohorts were used to determine the impact that different mTBI dosages had on genotype specific circadian/sleep profiles. An emphasis was placed on behavioral defects that occurred during the critical nighttime period (CT12-24) that highlights key features indicative of sleep fragmentation [13,29,36]. Several genotypes, including *Ref(2)P^c/e^* homozygous nulls (Supplemental **Figure S2**), *Atg8a^1^* and *Atg8a^2^* mutants (data not shown) demonstrated significant baseline activity and circadian cycle defects, even without trauma exposure. With a single mTBI (1x) bout, most flies did not survive for the entire time of the circadian study and were not included in the final analyses (**Figure S2**).

Young male fly cohorts were fully entrained in 12-hr light:dark conditions (**LD**) exposed to mTBI, allowed to recover and assayed signally using the DAM and data collection software and dark conditions (**DD**) [13]. Representative actograms (5 days, 30-min bins) of WT male flies (*w^1118^*/+) exposed to 0x, 1x, 5x, or 10x mTBI bouts demonstrated individual rhythmic activity and rest (inactivity) patterns, during the subjective daytime (**CT0-12**) and subjective nighttime periods (**CT12-24**, **Figure 4A**). For young WT male flies (*w^1118^/+*), a single mTBI bout had a modest but significant impact on several behavioral parameters, including daily sleep time (min/24-hrs/fly), activity (crossings/24-hrs/fly) and waking activity levels (crossings/min/fly, **Table 3**). However, a single mTBI bout did not significantly alter activity and sleep bout lengths (min/fly), sleep bouts (12-hr) or brief awakenings (12-hr) numbers that occurred during the key CT12-24 nighttime period (**Figure 4B**). As seen previously with females, outcrossed WT male flies (*w^1118^/+*) also showed a dose dependent alteration to circadian behaviors following exposure to different traumatic injury bout numbers (5x to 10x) [13]. This included activity and sleep related defects, including altered rhythmicity and daily activity patterns as well as increased sleep fragmentation levels (**Table 3**, **Figure 4**).

**Figure 4.**
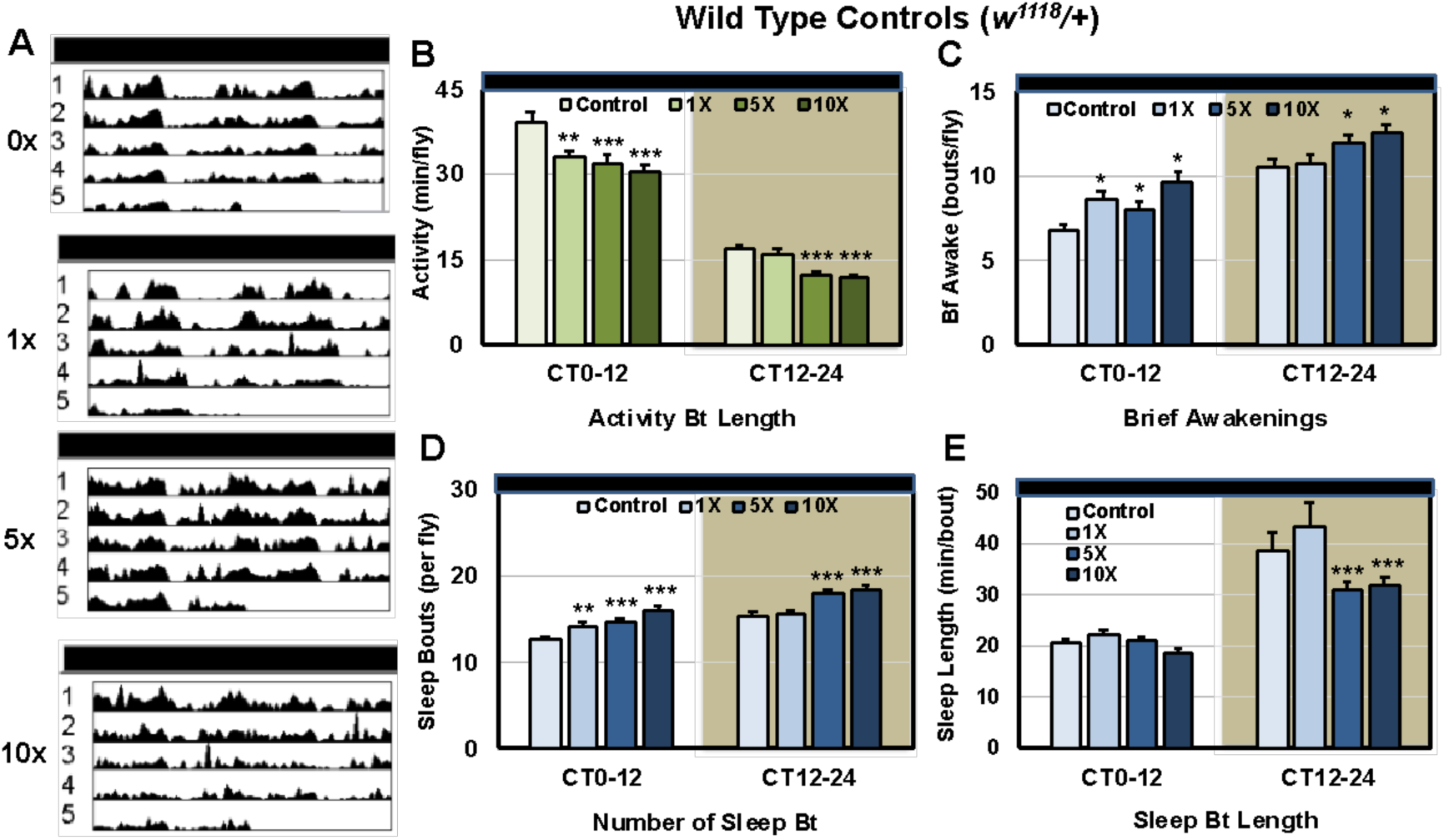
Dose dependent changes to control circadian-sleep behaviors following TBI. WT male fly cohorts were exposed to 0, 1x, 5x or 10x bouts of mTBI (2.1 m/s), allowed to recover for 5-days before being placed into the DAM monitoring system and assessed in constant darkness (**DD**). **A**) Representative double-plotted actograms of mTBI treated flies. **B-E**) Using the MATLAB-based software, analysis of sleep-related behaviors was performed to assess behaviors during subjective 12-hr day (CT0-12) and 12-hr night (CT12-24) time periods (n=32 each). **B**) Activity bout length (mins/12-hrs/fly), **C**) brief awakenings (no./12-hrs/fly), **D**) sleep bouts (no./12-hrs/fly) and **E**) sleep bout length (mins/12-hrs/fly). P values **≤ 0.01; ***≤ 0.001.

**Table 3.**
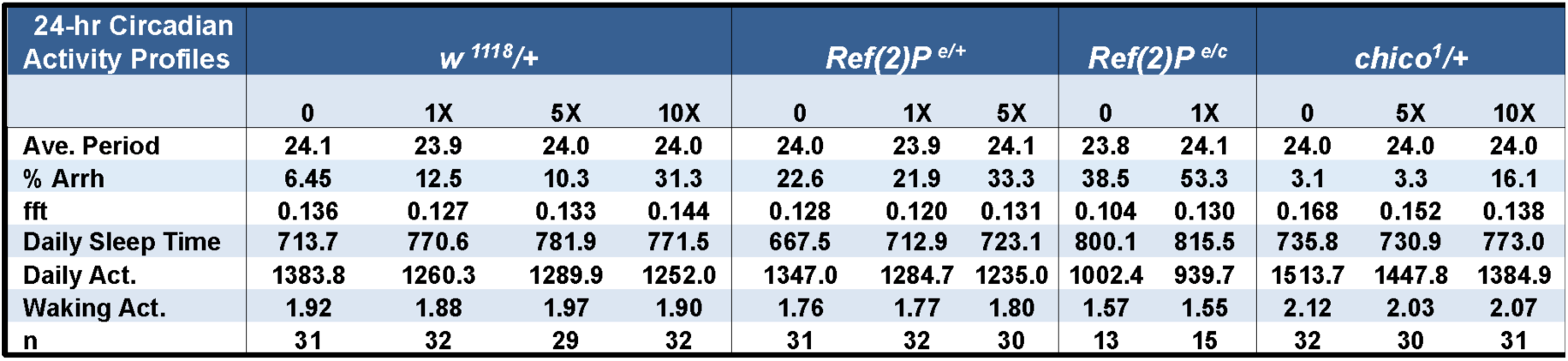
24-hour Circadian Profiles following Trauma (Sleep Lab Analysis)

Using these modified techniques, we examined the circadian/sleep behavioral profiles of young male flies from potentially trauma resistant and sensitive backgrounds. The baseline NRG and circadian behaviors of young *Ref(2)P^e^*/+, *Ref(2)P^e/c^* and *chico^1^/+* mutants were relatively normal. Therefore, these genotypes were used for subsequent trauma-based behavioral studies. An notable exception was adult male *Ref(2)P^e/c^* flies, which had robust basal NGR responses (**Fig. X**), but showed significant defects in circadian-sleep behavior profiles. Without being exposed to trauma, the *Ref(2)P^e/c^* flies died quickly once placed in the small assay tubes and DAM monitors (**Table 3**, **Supplemental Figure S2, Table S1**) [26]. The high mortality and arrhythmic rates of *Ref(2)P^e/c^* flies further indicated this genotype’s sensitivity to stress and did not facilitate multi-day assays using the DAM system. Indeed, the rhythmic, sleep and activity profiles of *Ref(2)P^e/c^* flies were largely unresponsive following 1x or 5x mTBI bouts (**Table 3**, **Supplemental Figure S2, Table S1**) [13].

The heterozygous *Ref(2)P^e^*/+ and *chico^1^/+* heterozygous flies overall showed relatively normal basal circadian, activity and sleep profiles, which permitted the assessment of trauma responses (**Table 3**). After exposure to a single mTBI (1x) bout, *Ref(2)P^e^*/+ mutants flies showed significant changes to activity and sleep-based parameters, especially during the key CT12-24 time period (**Table 3**; **Figure 6**). In contrast, the *chico^1^/+* mutants maintained robust rhythmicity, sleep and activity behavior profiles during the CT12-24 time-period that was largely unaffected even after high mTBI exposure levels (5x to 10x, **Supplementary Table S1 Table S1**). Only after *chico^1^/+* flies were exposed to the highest mTBI conditions (10x, **Table 3**, **Figure X**) was a significant change to activity and sleep-related profiles detected for this genotype [31]. Overall, the different behavior responses seen between WT (*w^1118^/+*), *Ref(2)P^e^*/+ and *chico^1^/+* fly genotypes following mTBI were generally consistent with other assays showing more global trauma responses, including severe MI^24hr^ and lifespan profiles. Taken together, we have developed an integrated assessment platform that can functionally identify genetic modifiers that show either resistance or sensitivity to whole body trauma and brain injury [55].

**Figure 5.**
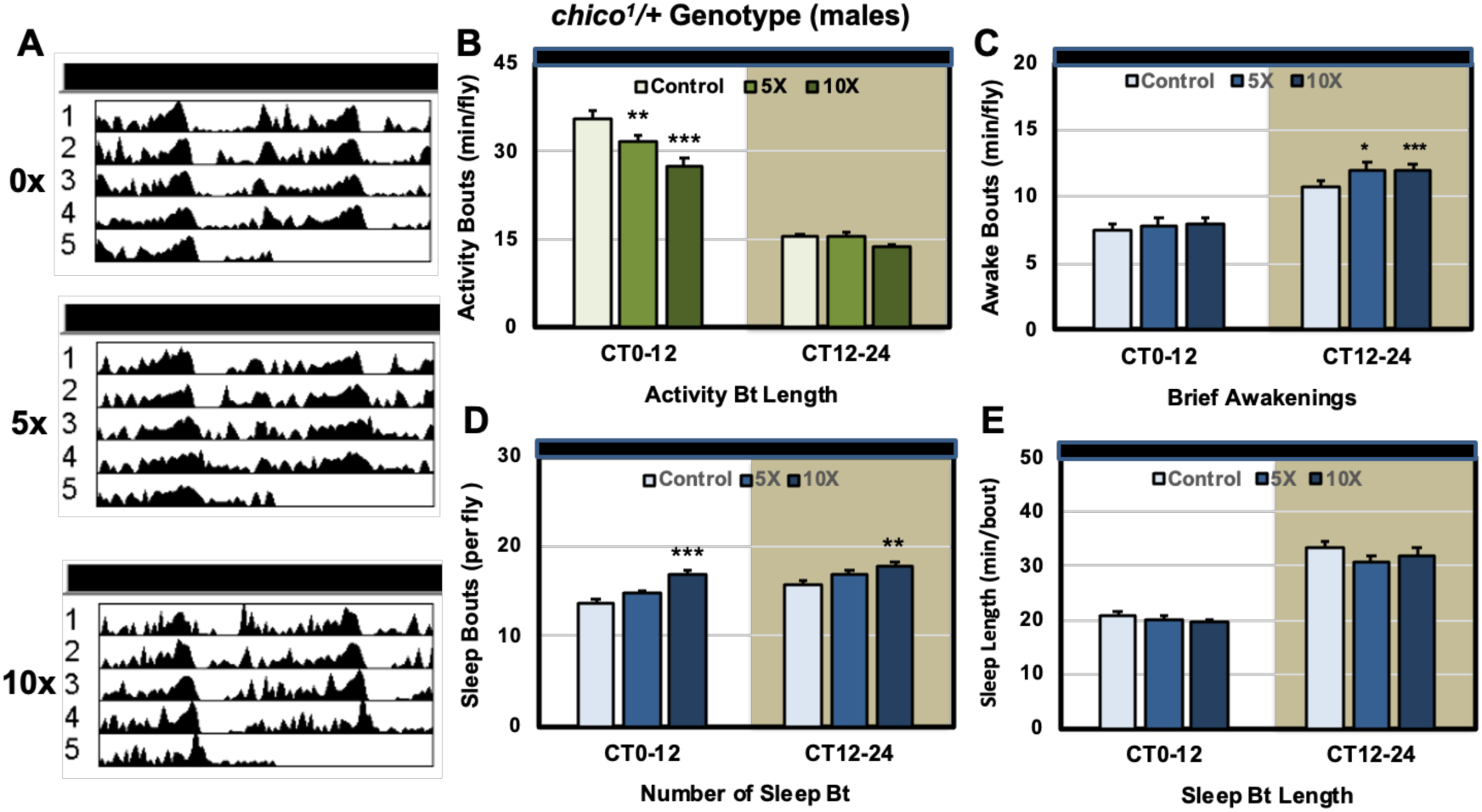
Dose dependent changes to *chico^1^/+* circadian-sleep behaviors following mTBI. *chico^1^/+* males exposed to 0, 5x or 10x mTBI bouts. A) Representative double-plotted actograms for control and mTBI treated flies. **B-E**) Subjective 12-hr day (CT0-12) and 12-hr night (CT12-24) time-period values (n=32). **B**) Activity bout length (mins/12-hrs/fly), **C**) brief awakenings (no./12-hrs/fly), **D**) sleep bouts (no./12-hrs/fly) and **E**) sleep bout length (mins/12-hrs/fly). P values * ≤ 0.05; ** ≤ 0.01; *** ≤ 0.001.

**Figure 6.**
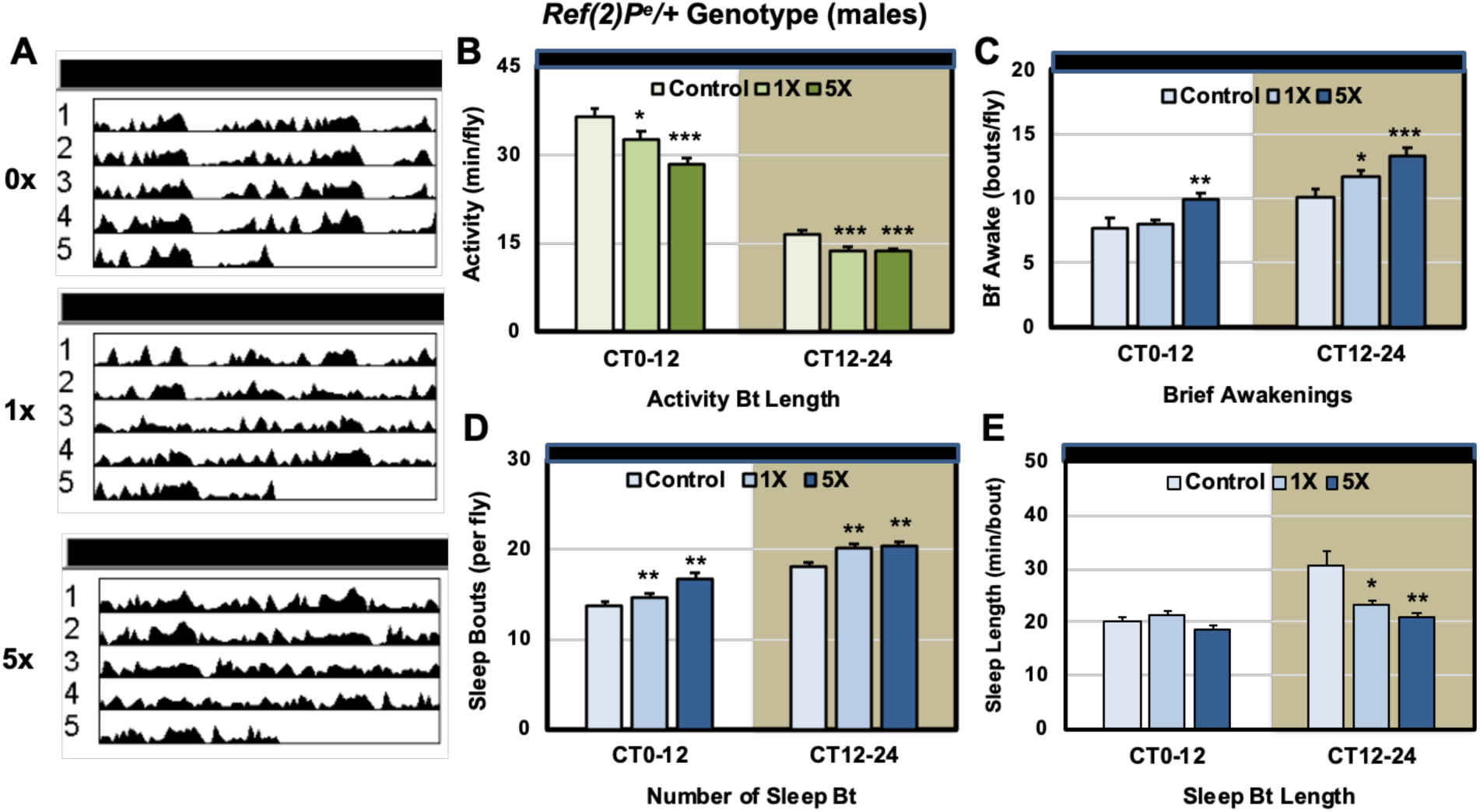
Dose dependent changes to *Ref(2)P^e^/+* circadian-sleep behaviors following mTBI. *Ref(2)P^e^/+* males were exposed to 0, 1x, 5x or 10x mTBI bouts. **A**) Double-plotted actograms for young control and mTBI treated cohorts. **B-E**) Subjective 12-hr day (CT0-12) and 12-hr nighttime (CT12-24) period values (n=32). **B**) Activity bout length (mins/12-hrs/fly), **C**) brief awakenings (no./12-hrs/fly), **D**) sleep bouts (no./12-hrs/fly) and **E**) sleep bout length (mins/12-hrs/fly). P values * ≤ 0.05; ** ≤ 0.01; *** ≤ 0.001.

## DISCUSSION

It’s been established that neural trauma exposure results in a complex series of responses, which in affected tissues that can be classified into “injury” categories [48,56]. This includes the primary injury resulting from direct and indirect mechanical forces that includes shearing, severing or crushing extracellular connections and structures as well as damaging intracellular structures and organelles [56]. In turn, trauma results in acute ROS production and the rapid upregulation of stress related factors and pathways including inflammation within neurons [5,48,56]. Often there is the coordinated activation of downstream signaling cascades that can facilitate the clearance of damaged structures and activate repair mechanisms [9,57,58]. Secondary injury can result from excessive or protracted inflammation and ROS production, that further exacerbates cellular damage and potentially slows repair of complex neural and glial structures and eventual functional recovery [9,57,58]. In humans, factors that further complicate our understanding of neural trauma is the relative inaccessibility of human neural tissues as well as complex cell populations, structures and interactions. Therefore, a rapid genetic model system that integrates multiple dynamic elements, ranging from age, gender to genetic differences, could provide unique insights into factors impacting both acute and long-term trauma responses and long-term recovery.

### Trauma Exposure and Mortality-Longevity Profiles

In this report, we examined the impact that exposure to severe or mild repetitive injury conditions had on adult *Drosophila* from several different genetic backgrounds. The tissue homogenizer system used for these studies produces uniform kinetic levels of closed traumatic injury (TBI) to the fly nervous system [13,29,49]. Individual TBI conditions were previously defined and established the dose-dependent decline to longevity and behavior profiles. The activation of neuronal autophagy and dynamic changes to *Drosophila* to inflammatory pathway components (AMPs) have also been optimized for this type concussive injury system [13,29,49]. Mammalian studies have also confirmed that TBI and SCI (spinal cord injury) exposure also results in autophagosome formation and accumulation, correlating with secondary injury responses [10,59,60,33,43]. This indicates that activating the autophagy pathway and bulk clearance is a conserved mechanism facilitating the removal and recycling of damaged intracellular components [24]. Therefore, the initial focus of this study was to compare the impact that gender and aging plays on global responses. Subsequent *in vivo* genetic studies determine whether modest changes to autophagy capacity (+/-) in neural tissues has on acute and long-term trauma responses, in terms of longevity, neural phenotypes and inflammation.

### The Impact of Age and Gender on *Drosophila* TBI Responses

A review of human TBI research highlights a substantial male bias, with men 40% more likely to experience reported head trauma than women [61]. However, these numbers are likely inaccurate since estimates from domestic violence survivors indicated that as high as 75% of individuals have sustained head injuries [17,61-63]. Patient data and rodent TBI studies further underscore the complexity that gender, hormonal differences, and chromosomal dosage plays on global trauma responses. Here, young wild-type male and female flies (*w^1118^*/+) showed similar acute mortality profiles following trauma (MI^24hr^, sTBI 1x), which increased sharply for both with age (**Table 1**). In concordance with our previous work, the age-dependent increase in sensitivity was consistent with the decline in neural autophagic capacity and an increase in the age-dependent stress sensitivity in older flies (**Table 1**) [25,31]. However, by 5-weeks female flies have lower MI profiles compared to age-matched male cohorts, likely reflecting the longer average lifespans of females. Overall, the *Drosophila* sTBI MI-24h profiles are similar with the relatively poor trauma outcomes experienced by elderly human patients [10,16]. In addition, the modest trauma resilience of aged female flies, is consistent their increased longevity and comparatively elevated levels of neural autophagy occurring later in life [9,27,31].

### The Impact of Genetic Backgrounds on Trauma Responses

The acute mortality (MI^24hr^, sTBI 1x) and longevity (mTBI, 10x) profiles were obtained from stress sensitive (*Ref(2)P*, *Atg8a*) or resistant (*chico*) genetic backgrounds. (**Tables 1-2**) [25,27]. Overall, the MI-24hr responses of *Atg8a^1^*and *chico*^1^/+ mutant adults were consistent with starvation or H_2_O_2_ exposure-based studies (**Tables 1, 2**) [25]. With sTBI the *Atg8a^1^* (reduced autophagy) and *chico*^1^/+ (enhanced autophagy) adult cohorts clearly showed expected sensitive and resistant trauma phenotypes, while adult *Ref(2)P* mutants stress responses hadn’t been examined in detail [26]. The Ref(2)P and p62/SQSTM1 family of proteins have clear roles within the autophagic pathway and intersect with the immune/inflammatory system pathways (NFκB, NRF2, cGAS/Sting) [26,38,64-66]. Further, human genetic studies have linked p62/SQSTM1 mutations with inherited neurological disorders [38,65,67,68]. We anticipated that loss of Ref(2)P protein would also increase adult sensitivity to trauma, which would include a further reduction to lifespans following exposure to severe and mild repetitive trauma (**Tables 1-2**) [24,26,69]. Imaging and Western blot analysis also confirm that *Ref(2)P^e/e^* mutants have attenuated neural autophagic levels that include reduced autophagosome formation and pathway flux (**Fig. 1**). The inability of *Ref(2)P* mutants to full activate neuronal autophagy is consistent with the overall trauma sensitivity of both homozygous and heterozygous adult mutants and has potential implications for people appear to have normal neural function but carry heterozygous SQSTM1 mutant combinations [51,67,70,71].

### Inflammation and TBI outcomes

Human patients that have relatively poor TBI outcomes, often demonstrate elevated or protracted inflammatory responses [4,55-57]. Inflammation is an essential injury signal that activates and recruits immune cells (macrophages) and/or microglia to injury sites [57,58]. Subsequent cellular events facilitate the clearance of damage and promote activation of remodeling and repair pathways [19]. However, when there is protracted or excessive levels of inflammation, often there are secondary rounds of tissue damage [2,6,18]. From neural therapeutics studies on neural aging and trauma phenotypes, basal and acute changes to *Drosophila* inflammatory markers or antimicrobial peptides (AMPs) following trauma were examined [9,26,28]. This included a range of treatments with CBD to active (LLG) and inactive (ReseT®) bacterial based biologics to injured fly cohorts [26,28]. Overall, treatments that promoted longevity, preserved behaviors and lowered neural aggregates in aged flies, while also having a positive effect on trauma responses [26,28]. Neural protective treatments also altered neural AMP expression levels and adult *Drosophila* inflammatory responses. Of the three distinct fly genotypes each had unique AMP profiles (**Figure 2**). The TBI sensitive *Ref(2)P^E^*/+ adults showed suppressed *DptB* levels before and after TBI (4-, 24-hrs. **Figure 2**). When compared to controls, resistant *chico^1^/+* adults started with control comparable basal AMP profiles but had a rapid and robust AMP activation (4-hrs post) that rapidly resolves after 24-hrs following trauma (**Figure 2**). This indicates that inflammatory responses in neural tissues correlate with global trauma response trends (+/-) for different genetic backgrounds. Further implications are that relatively silent genetic combinations, that don’t create obvious developmental defects or alter baseline physiology, could have a significant impact on stress related responses in humans.

### Behavioral Profiles as Phenotypic Markers of Trauma Response

Along with neuroinflammation, head trauma patients often show signs of altered neurological function. Acute problems can range locomotor defects, to sleep and circadian cycle dysregulation [47,48,59]. Previous therapeutic studies showed trauma related locomotor defects (geotaxis) in flies could be improved, indicated the negative geotaxis response assay is a robust assessment that can be used to detect subtle *in vivo* neurological changes [26,28]. Here we found the climbing response of young controls (1-week, *w^1118^*/+) and *chico^1^/+* mutants (1-, 3-weeks), while impacted, on averages well above the 25% decrease cutoff, indicating relative trauma resistance (**Figure 3**). Conversely, aged cohorts (3-week, *w^1118^*/+) or young autophagy mutants (*Atg8a^1^*, *Ref(2)P^c/e^*) hed more than a 25% or greater decrease in locomotor profiles following trauma, consistent with TBI sensitivety (**Figure 3**). Further, using the Gal4/UAS system to increase (OE) or reduce (KD) the *Ref(2)P* gene in different cell types, climbing studies indicated that elevating Ref(2)P in glial cells was linked to improved locomotor responses following mTBI (**Figure 3**).

Another common problem seen with human trauma patients are substantial alterations to sleep patterns, including insomnia, hypersomnia and sleep fragmentation [48,49,59]. To further characterized TBI resistant or sensitive genotypes, sleep related behaviors with different levels of trauma were directly examining. Initially, adult male flies from sensitive (*Atg8a^1^*, *Ref(2)P^e/c^, Ref(2)P^e^/+*) and resistant (*chico^1^/+*) genotypes were examined (**Figures 4-6**). However, homozygous *Atg8a^1^* and *Ref(2)P^e/c^* mutant cohorts showed base-line circadian defects and with trauma died quickly, preventing their full analysis using the DAM assay system (**Supplemental Figure 2**) [9,26]. Indeed, to permit the dose-dependent analysis of sleep-circadian patterns, the sensitive *Ref(2)P^e^/+* flies required lower trauma levels to be used (1x to 5x mTBI bouts). Consistent with other TBI responses, the *chico^1^/+* males easily survived the assays and showed minimal disruption of activity levels, rhythmicity, and sleep patterns, even after exposure to mTBI (10x) (**Figure 5**). Overall, the key marker of dysregulation in flies or sleep fragmentation, was not significantly altered during the critical CT12-24 (dark) period (**Figure 5**). Conversely, a single mTBI (1x) bout disrupted sleep and activity patterns of *Ref(2)P^e^/+* heterozygous males, which was exacerbated further with 5x bouts and caused extensive lethality after receiving 10x bouts (**Figure 6**). Taken together with other physiological responses, a consistent pattern emerges that adult *Drosophila* trauma models are an effective approach to define the relative impact that gender, age and potential genetic factors may have on both acute responses and long-term trauma outcomes. These models have the potential to help build on the key molecular mechanisms involved with signaling, repair and remodeling systems influencing functionality after trauma.

### Implications for Human Health

There are several takeaways or implications from our Drosophila TBI studies with relationship to human health. The first is best highlighted by ongoing advances in next sequencing technology, detailed curation and unprecedented access to human biomedical genomics/genetics information. We used the Genome Aggregation Database (gnomAD, https://gnomad.broadinstitute.org/news/2023-11-gnomad-v4-0/) to target the human *Ref(2)P* homologue, for human Sequestosome-1 (SQSTM1) mutations. Initially, SQSTM1 defects were associated with familial cases of Paget disease of bone but now there are significate links to amyotrophic lateral sclerosis (**ALS**) [30,45,65,66]. Exome sequencing studies are examining loss-of-function SQSTM1 mutations and their potential linkage with other early onset neurodegenerative disorders [67,68]. The gnomAD browser was also used to review sequence variants and alleles for human versions of the *Atg8a* (MAP1LC3A, MAP1LC3B, MAP1LC3B2, MAP1LC3C, GABARAP), and *chico* (IRS1, IRS2) genes [60,61]. Human *Atg8a* orthologs (GABARAPs and MAP1LC3s) contain several genetic variants, however, none are predicted to significantly alter the genes expression patterns or result in functional protein changes. Nor are they linked to genetic disorders, which is further highlighted by negative loss-of-function observed/expected upper bound fraction (LOEUF) scores for each gene. The implications are the full function of these genes are critical for multiple developmental and physiological processes, resulting in the intolerance for any type of genetic variation [21,24]. Indeed, the two *Atg8a* fly mutations used in this study are P-element promoter insertions that lowers global expression and are not represented in human sequences as altered alleles.

GnomAD assessments of the IRS1 and IRS2 genes identifies several allelic variants with positive LOEUF scores in human populations, with several associate with an increased risk of Type-II diabetes and lipodystrophy [62-64]. A direct genetic association of IRS1/IRS2 defects with neural degenerative disorders or trauma sensitivity haven’t been identified. However, both dietary and genetic-based studies clearly show that activated insulin pathway signaling is a potent suppressor of autophagic function in multiple tissue types. The TBI resistance of *chico^1^*/+ fly cohorts is consistent with overall positive effect of suppressed insulin signaling and elevated autophagic capacity, which in turn promotes neural stress resistance and longevity [24,27]. The potential for high calorie Western diets, obesity and Type-II diabetes to be significant non-genetic risk factors for poor head trauma outcomes even at relatively young people in also raised. The overall implications are that decreased autophagic capacity in neurons, whether from excess calories, dysregulated insulin signaling or modest genetic defects to the autophagy pathway, will likely have a significant impact on the outcome of trauma patients.

## Abbreviations

TBI: traumatic brain injury
mTBI: mild repetitive TBI
sTBI: severe TBI
Genome-wide association studies: GWAS
Ref(2)P: Refractory to sigma P
SQSTM1: Sequestosome-1
AD: Alzheimer disease
ATG5: autophagy related 5
ATG7: autophagy related 7
ATG12: autophagy related 12
BECN1: beclin 1, autophagy related
CCI: controlled cortical impact
MAP1LC3/LC3: microtuble-associated protein
PE: diacyl glycerophosphatidylethanolamine
ROS: reactive oxygen species

## Acknowledgements

This work was supported by R01AG039628 from NIH/NIA (KDF).

## Author Contributions

K.D.F. and E.P.R. conceived of and directed the experiments. B.M., N.E.M., K.D.F., and E.P.R. conducted the experiments. B.M., J.J.R., M.M.L., K.D.F., and E.P.R. analyzed the results. J.J.R., M.M.L., K.D.F., and E.P.R. prepared the manuscript. All authors reviewed the manuscript.

## Additional Information

