## Supplemental Data for "Genetic Modifiers Influencing the Acute and Long-Term Responses to Traumatic Brain injury in *Drosophila*"

**Supplemental Figure S1.**

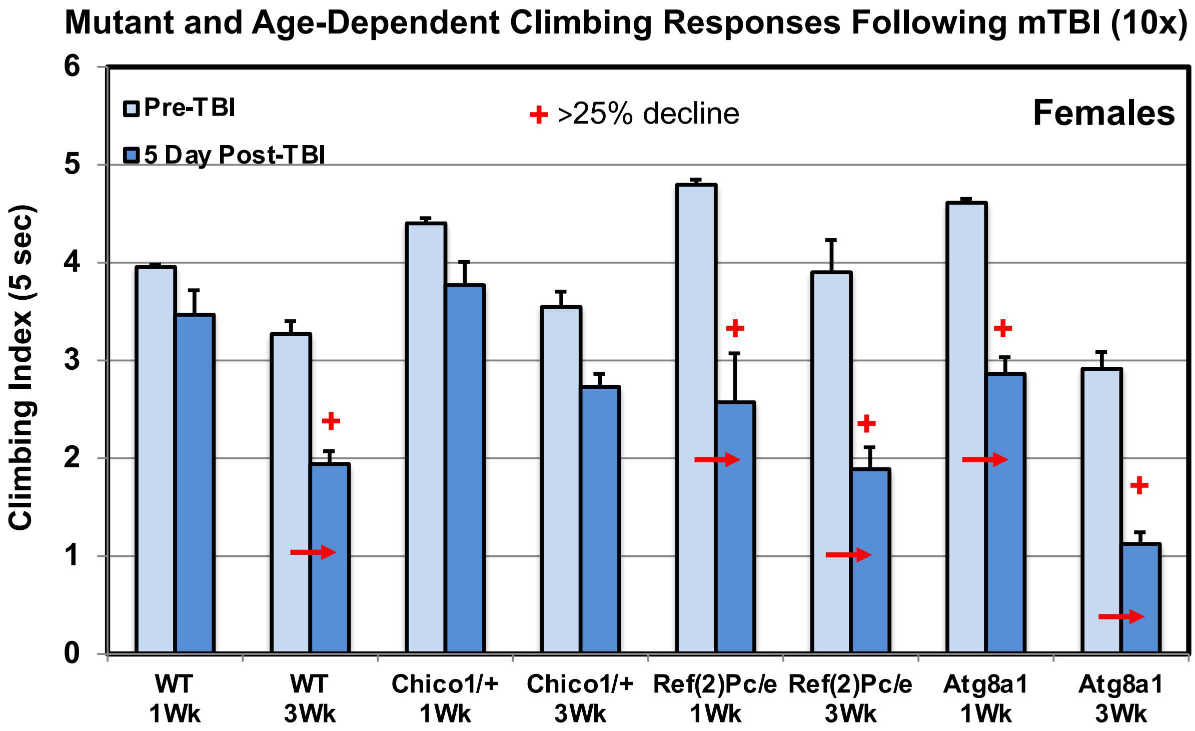


**Supplemental Figure S1. Genotype and Age-Related Changes to Climbing Behaviors (NGR) of Adult Female *Drosophila* Before and After mTBI (10x).** The NGR profiles of 1 or 3-week old WT ($w^{1118}$/+), $chico^{1}$/+, $Ref(2)P^{e/c}$ null and $Atg8a^{1}$ mutant fly cohorts before and 5-days following mTBI (10x). A decrease in climbing behavior profiles greater than 22% before and after trauma exposure are highlighted (+).

**Supplemental Figure S2.**

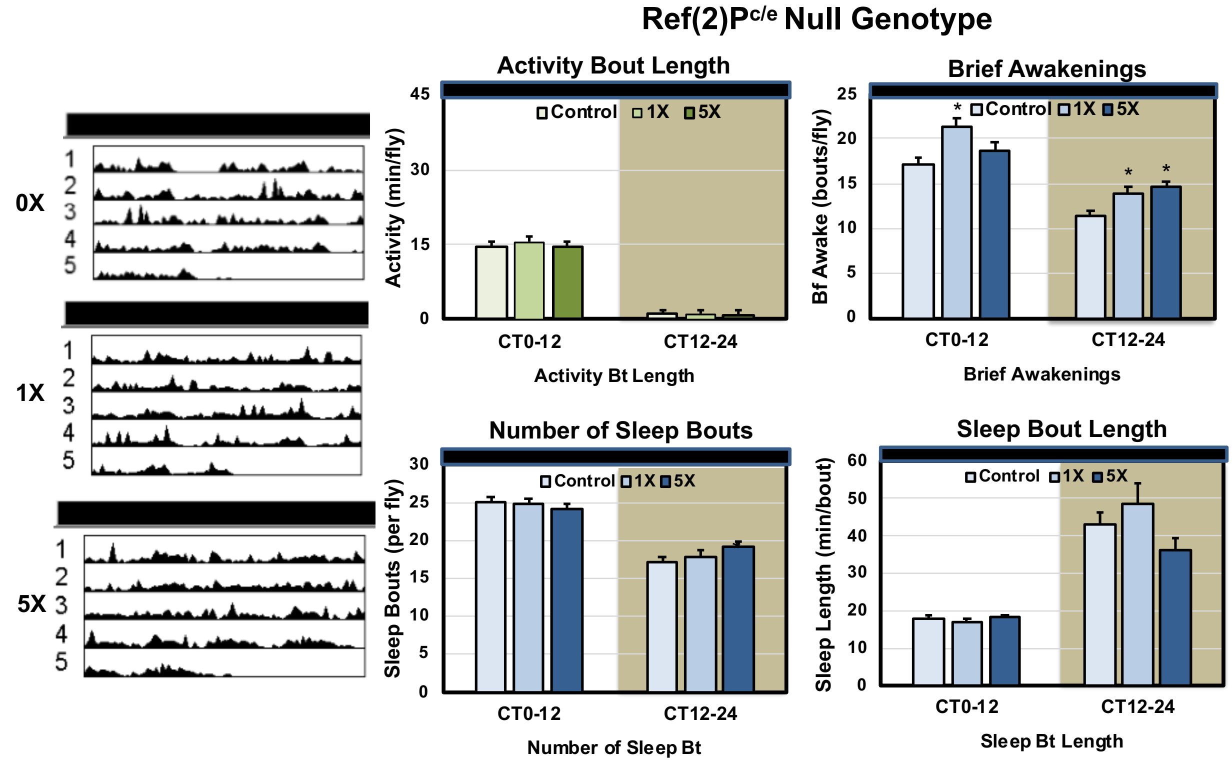


**Supplemental Figure S2. Dose dependent changes to Ref(2)P circadian-sleep behaviors following TBI.** WT male fly cohorts were exposed to 0, 1x, 5x or 10x bouts of mTBI (2.1 m/s), allowed to recover for 5-days before being placed into the DAM monitoring system and assessed in constant darkness (**DD**). **A**) Representative double-plotted actograms of mTBI treated flies. **B-E**) Using the MATLAB-based software, analysis of sleep-related behaviors was performed to assess behaviors during subjective 12-hr day (CT0-12) and 12-hr night (CT12-24) time periods (n=32 each). **B**) Activity bout length (mins/12-hrs/fly), **C**) brief awakenings (no./12-hrs/fly), **D**) sleep bouts (no./12-hrs/fly) and **E**) sleep bout length (mins/12-hrs/fly). P values **≤ 0.01; ***≤ 0.001.

### Supplemental Table S2. Sleep Lab CT0-CT12 and CT12-CT24 Behavior Profiles.

| 12-hr LD Profiles (Ave per fly) | | *w* $^{1118}$/+ | | | | *Ref(2)P* $^{e/+}$ | | | *Ref(2)P* $^{e/c}$ | | *chico*$^{1}$/+ | | |
|---|---|---|---|---|---|---|---|---|---|---|---|---|---|
| | | 0 | 1X | 5X | 10X | 0 | 1X | 5X | 0 | 1X | 0 | 5X | 10X |
| Brief Awakenings | CT0-12 | 6.77 | 8.58 | 8.05 | 9.69 | 7.75 | 7.99 | 9.87 | 16.86 | 15.52 | 7.49 | 7.86 | 8.01 |
| | CT12-24 | 10.50 | 10.76 | 12.04 | 12.68 | 10.12 | 11.68 | 13.31 | 15.29 | 14.91 | 10.68 | 11.93 | 11.92 |
| Active Time | CT0-12 | 465.3 | 424.2 | 427.9 | 434.6 | 460.3 | 432.9 | 426.2 | 367.1 | 362.3 | 447.3 | 433.1 | 408.7 |
| | CT12-24 | 261.2 | 245.3 | 230.4 | 233.9 | 312.2 | 294.5 | 290.8 | 273.0 | 262.3 | 257.1 | 276.2 | 258.5 |
| Sleep Time | CT0-12 | 254.7 | 295.8 | 292.1 | 285.4 | 259.7 | 287.1 | 293.8 | 352.9 | 357.7 | 272.7 | 286.9 | 311.3 |
| | CT12-24 | 458.8 | 474.7 | 489.6 | 486.1 | 407.8 | 425.5 | 429.2 | 447.0 | 457.7 | 462.9 | 443.8 | 461.5 |
| Activity Bout Length | CT0-12 | 39.4 | 33.1 | 32.0 | 30.5 | 36.5 | 32.6 | 28.4 | 20.6 | 21.3 | 35.6 | 31.8 | 27.6 |
| | CT12-24 | 16.9 | 15.9 | 12.4 | 11.9 | 16.6 | 13.8 | 13.6 | 13.4 | 14.7 | 15.4 | 15.6 | 13.8 |
| Sleep Bout Length | CT0-12 | 20.8 | 22.4 | 21.1 | 18.8 | 20.3 | 21.5 | 18.4 | 18.7 | 18.0 | 20.8 | 20.1 | 19.6 |
| | CT12-24 | 38.7 | 43.5 | 31.3 | 31.8 | 30.5 | 23.2 | 29.1 | 28.6 | 28.4 | 33.3 | 30.5 | 31.9 |
| Sleep Bout Number | CT0-12 | 12.7 | 14.3 | 14.8 | 16.1 | 13.8 | 14.7 | 16.8 | 21.5 | 20.5 | 13.7 | 14.7 | 16.8 |
| | CT12-24 | 15.5 | 15.7 | 18.1 | 18.5 | 18.0 | 20.1 | 20.3 | 20.7 | 19.7 | 15.7 | 16.8 | 17.7 |

One-way ANOVA followed by Turkey's multiple comparison post-hoc test. For data regarding multiple time points, results were compared by two-way repeated measures ANOVA, followed by Bonferroni's multiple comparison test. Cross symbols represent significance when comparing 3 groups in ANOVA, excluding the unmarked treatment group from the analysis. Data averages were and statistical analyses between groups and were calculated using the feature of GraphPad All values are reported as means + SEM.
